# Anti-tumor effects of antimicrobial peptides, targets of the innate immune system, against hematopoietic tumors in *Drosophila mxc* mutants

**DOI:** 10.1101/452961

**Authors:** Mayo Araki, Rie Awane, Tetsuya Sato, Yasuyuki Ohkawa, Yoshihiro H. Inoue

## Abstract

The innate immune response is the first line of defense against microbial infections. In *Drosophila*, three immune pathways induce the synthesis of antimicrobial peptides (AMPs) in the fat body. Recently, it has been reported that certain cationic AMPs exhibit selective cytotoxicity against human cancer cells. However, little is known about their anti-tumor effects. *Drosophila mxc^mbn1^* mutants exhibit malignant hyperplasia in a larval hematopoietic organ called the lymph gland (LG). Here, using RNA-Seq analysis, we found that many immunoresponsive genes, including AMP genes, were up-regulated in the mutants. Down-regulation of these pathways by either a *Toll* or an *imd* mutation enhanced the tumor phenotype of the *mxc* mutants. Conversely, ectopic expression of each of five different AMPs in the fat body significantly suppressed the LG hyperplasia phenotype in the mutants. Thus, we propose that the *Drosophila* innate immune system can suppress progression of hematopoietic tumors by inducing AMP gene expression. Overexpression of any one of these five AMPs resulted in enhanced apoptosis in the mutant LGs, while no apoptosis signals were detected in controls. We observed that two AMPs, Drosomycin and Defensin, were taken up by circulating hemocyte-like cells, which were associated with LG regions showing reduced cell-to-cell adhesion in the mutants; another AMP, diptericin, was directly localized on the tumors without intermediating hemocytes. These results lead us to conclude that the AMPs have a specific cytotoxic effect that enhance apoptosis exclusively in the tumor cells.

**Summary statement:** Antimicrobial peptides can be associated with tumor cells generated in a hematopoietic tissue in *Drosophila mxc* mutants and have an anti-tumor effect in suppressing their growth.

## Introduction

To combat invading microbial pathogens with which they share the same environment, multicellular organisms have developed robust defense mechanisms. Insects rely fundamentally on multifaceted innate immunity, involving both humoral and cellular defense responses, as they lack an acquired immunity system (Brennan and Anderson, 2004; Hoffmann, 2003; Tzou et al., 2002). The hallmark of the humoral reactions is the systemic antimicrobial response. This involves the challenge-induced synthesis of antimicrobial peptides (AMPs) in the fat body, which is a major immune-responsive tissue as well as an approximate functional equivalent of the mammalian liver. AMPs secreted into the hemolymph destroy invading microorganisms (Lemaitre and Hoffmann, 2007; Zhang and Gallo, 2016). To date, seven distinct AMPs plus their isoforms have been identified in *Drosophila melanogaster*: drosomycin, defensin, diptericin, metchnikowin, attacin-A, cecropin A2 and drosocin (Lemaitre and Hoffmann, 2007). A series of genetic analyses have demonstrated that the genes encoding these AMPs are regulated by three kinds of innate immune signaling pathways (Hoffmann and Reichhart, 2002; Tzou et al., 2002). First, the Toll-mediated pathway is activated mainly by fungi and Gram‐positive bacteria (Lemaitre et al., 1997). Infection by these microbes initiates activation of serine protease cascades, which ultimately lead to the production of the active ligand (Jang et al., 2006). The activated ligand protein binds directly to the transmembrane receptor Toll and activates it (Morisato and Anderson, 1994). Activated Toll transmits a signal through several proteins in the cytoplasm. At the end, the protein kinase Pelle phosphorylates and decomposes Cactus, a cytoplasmic ankyrin repeat-containing protein. In the absence of infection, Cactus prevents the NF-_κ_B family transcription factors Dorsal (Dl) and Dorsal-related immune factor (Dif) from entering the nucleus. Once Cactus is decomposed in response to the Toll signal, Dl and Dif are free to translocate into the nucleus, where they can induce the expression of antimicrobial peptide genes such as *Drosomycin* (*Drs*) (Belvin and Anderson, 1996; Brennan and Anderson, 2004; Fehlbaum et al., 1994; Hoffmann, 2003). Second, the Imd-mediated pathway responds mainly to infection by Gram‐negative bacteria. Cell wall components of the bacteria are recognized by specific transmembrane receptors (Choe et al., 2002; Gottar et al., 2002; Lim et al., 2006; Takehana et al., 2004). This leads to activation of the signaling pathway mediated by a complex composed of multiple proteins including the adaptor protein Imd (Choe et al., 2005). Subsequently, the protein complex phosphorylates the Relish transcription factor. This phosphorylation triggers the cleavage of Relish (Ertürk-Hasdemir et al., 2009; Stöven et al., 2000; Tanji and Ip, 2005). The N-terminal NF-_κ_B modules of the transcription factor drive expression of genes encoding antimicrobial peptides, such as *Diptericin* (*Dip*). Third, JAK-STAT is also involved in the innate immunity pathway (Dostert et al., 2005). Activation of the JAK-STAT pathway commences when a ligand called Upd binds to its receptor, Domeless (Dome). This allows the association of JAK/Hop with the receptor. Activated Hop in turn phosphorylates Dome, resulting in the formation of docking sites for the cytoplasmic STATs. The recruitment of STATs induces STAT phosphorylation and ultimately transcription of the *Turandot* genes (An et al., 2012; Arbouzova and Zeidler, 2006). The Toll-mediated, Imd-mediated and JAK-STAT pathways are equivalent to the mammalian TLR-, TNFR‐ and JAK/STAT-mediated signaling pathways, respectively (Hoffmann, 2003; Rawlings et al., 2004; Valanne et al., 2011). Therefore, *D. melanogaster* has been regarded as a powerful model organism for the investigation of innate immune signaling pathways, especially humoral defense. A few studies have reported that the Toll-mediated and JAK-STAT signaling pathways are activated not only by invading microbes but also by tumor cells (Parisi et al., 2014; Pastor-Pareja et al., 2008). However, little is known about the mechanism by which an innate immune pathway is activated in the fat body of organisms harboring tumors.

In addition, cellular immune defenses also depend greatly on hemocyte-mediated phagocytosis by a macrophage-like cell called the plasmatocyte. In the case of larger invading microorganisms, specialized flattened cells called lamellocytes congregate around the intruders and encapsulate them (Lanot et al., 2001; Rizki and Rizki, 1980). Some evidence suggesting that *Drosophila* can respond to tumor cells in the body and invoke its immune system has been obtained. Once the basement membrane has been disrupted, circulating hemocytes increase in number and become associated with the tumors to restrict tumor growth (Pastor-Pareja et al., 2008).

In this study, we used a lethal mutant of the *multi sex combs* (*mxc*) gene, *mxc^mbn1^*, as a *Drosophila* hematopoietic tumor model (Sherestha and Gateff, 1982; Santamaria and Randsholt, 1995; Remillieux-Leschelle et al., 2002). *mxc^mbn1^* mutants exhibit hyperplasia in a larval hematopoietic tissue called the lymph gland (LG), whereas they show reduced tissue growth in the imaginal discs. The mutants consequently die at the larval/pupal stage, perhaps due to both of these effects. Mutant larvae contain increased numbers of circulating hemocytes and abnormally differentiated hemocytes in the hemolymph. LG cells isolated from the mutant larvae could further proliferate, invade host tissues and ultimately kill the host, when they were implanted into normal adult abdomens (Remillieux-Leschelle et al., 2002). In contrast, wild-type LG cells no longer proliferated in the host. Thus, the LG tumors in *mxc^mbn1^* can be considered as malignant tumors (Remillieux-Leschelle et al., 2002). The wild-type gene is also essential for development of the imaginal discs, as mutants for some *mxc* alleles showed defective disc growth and a lack of germ cells (Santamaria and Randsholt, 1995; Saget et al., 1998; Landais et al., 2014)

Here, we first present genetic evidence indicating that the innate immune pathways are activated in the *mxc^mbn1^* mutants carrying the LG tumors. Second, we show that this activation is essential for suppression of tumor progression in the hematopoietic tissue. Moreover, we demonstrate that the anti-tumor effect is exerted through induction of AMPs, which are targets of the innate immune pathways. We further observed that AMPs secreted into the hemolymph were preferentially associated with LG tumors showing a reduced cell-cell adhesion, by direct binding or with the aid of circulating hemocytes. Overexpression of the AMPs enhanced apoptosis in the LG of *mxc^mbn1^*, whereas no apoptosis occurred in control LGs. These results lead us to propose that a kind of surveillance system that activates the innate immune system in response to tumor cells exists in *Drosophila*. These AMPs may represent promising new anti-cancer substances without side-effects.

## Results

### Malignant blood neoplasm phenotype appears in larvae hemizygous for *mxc^mbn1^* mutation

Previous studies demonstrated that *mxc^mbn1^* hemizygotes showed hyperplasia of a larval hematopoietic tissue, the lymph gland (LG), in which precursors of hemocytes abnormally increased in number. The LG in wild-type larvae is arranged bilaterally so that its right and left halves flank the dorsal vessel at the midline (Figure 1A). Clusters of hematopoietic cells comprising a hemisphere of the gland are called lobes; these are aligned segmentally in pairs along the anterior-posterior axis, as a larger primary (1^st^) lobe, a secondary (2^nd^) lobe and a tertiary (3^rd^) lobe. Each lobe in the wild-type LG is distinctively separated by a pericardial cell (arrows and arrowheads, Figure 1B). Whole lobe regions of a right or left half of the LG preparations prepared from single larvae and stained with DAPI were 0.04 ± 0.004 mm^2^ on average (n = 22). By contrast, the LG from *mxc^mbn1^* mutants at a mature larval stage possessed enlarged lobes containing extra numbers of pre-hemocytes, thus exhibiting a LG tumor overgrowth (Figure 1C). Whole lobe regions of the mutant LGs were 0.08 ± 0.009 mm^2^ on average (n = 23), twice as large as those of wild-type (*p* < 0.0001, Student’s t test) (Figure 1D). Notably, the posterior lobes were more prominently enlarged, while the primary lobe was often undetectable or was dislocated, possibly as a consequence of a tissue burst derived from hyper-proliferation of the hematopoietic cells (Figure 1C). These mutant larval LG cells continue to proliferate and invade into other tissues when the mutant cells are injected into abdomens of normal adults (see Introduction). Thus, it is possible to assert that the *mxc^mbn1^* mutant exhibits a malignant phenotype.

**Figure 1.**
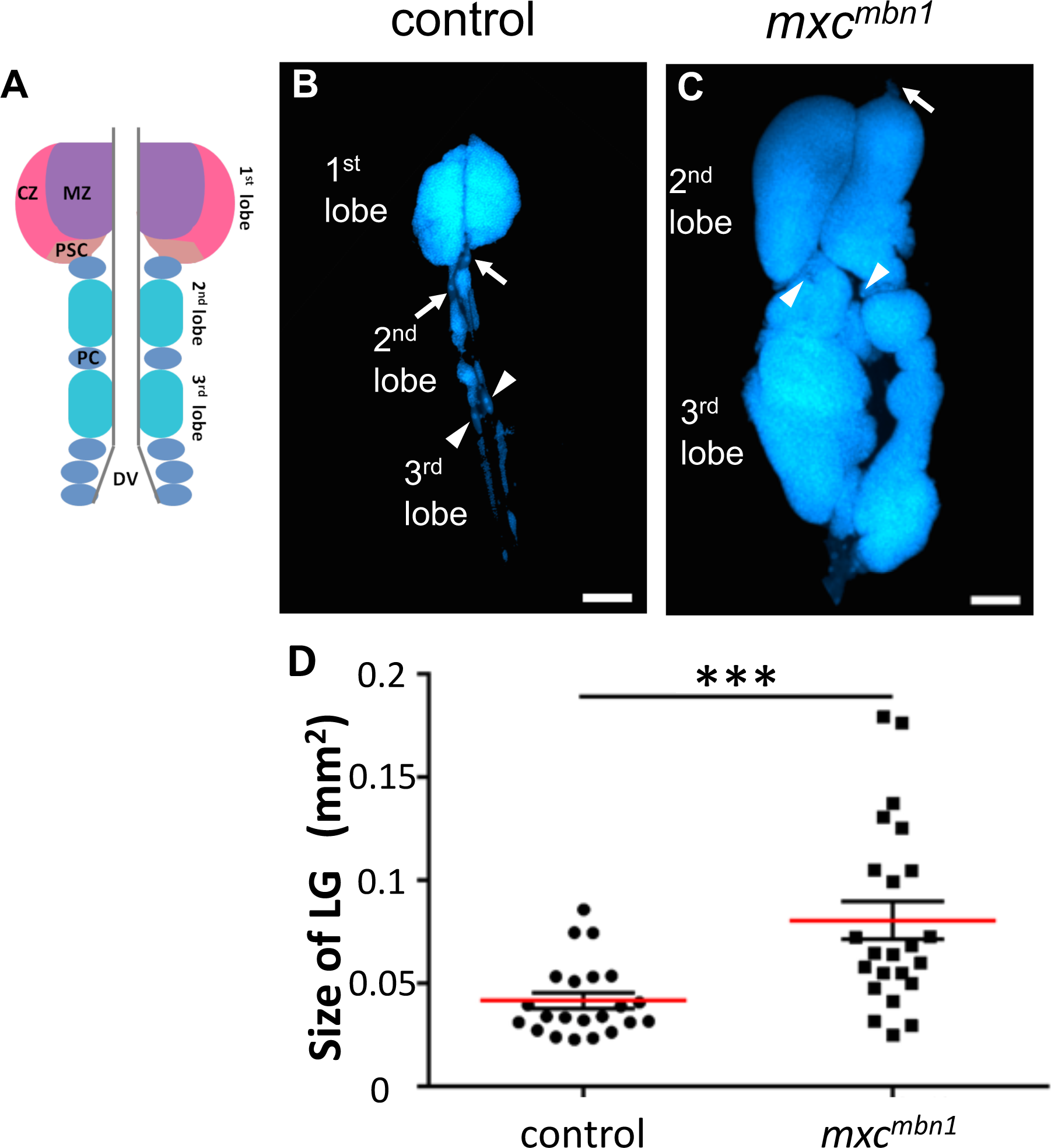
Hyperplasia of the larval lymph gland in *mxc^mbn1^* larvae. (A) Schematic representation of the LG in *Drosophila* 3^rd^ instar larvae. (B, C) DAPI-stained LGs isolated from wild-type (B) or hemizygous larvae (*mxc^mbn1^/Y*) for *mxc^mbn1^* (C). (B) The wild-type LG is arranged bilaterally so that it flanks the dorsal vessel at the midline. Each lobe is distinctively separated by a pericardial cell (arrows: pericardial cells separating the 1^st^ lobe from the 2^nd^ lobe; arrowheads: pericardial cells separating the 2^nd^ lobe from the 3^rd^ lobe). (C) LG tumor overgrowth in the *mxc^mbn1^*mutant larva. The posterior lobes are more prominently enlarged, while the 1^st^lobe, which was located anterior to the pericardial cell (arrow), has been lost in this LG. (Bars: 100 μm.) (D) Quantification of tumor size. The right or left halves of whole LG regions of the mutant LGs (0.08 mm^2^ on average, n > 20) are twice as large as those of wild-type (0.04 mm^2^ on average, n > 20) (Student’s t test, \*\*\**p* < 0.001. Error bars represent SEM. Red horizontal lines represent average.).

### RNA sequence analysis demonstrates that many immune response genes are up-regulated in *mxc^mbn1^* larvae

To identify genes whose mRNA levels were altered in the LG malignant hyperplasia in *mxc^mbn1^* larvae, we carried out RNA-Seq analysis. We determined the DNA sequences of 19,529,450 cDNA reads prepared from mRNAs expressed in wild-type mature larvae and 26,290,482 cDNA reads from *mxc^mbn1^* larvae at the same developmental stage. The RNA-Seq reads were mapped to a total of 13,747 mRNAs. We found 209 genes for which the mRNA levels in the mutant increased to more than ten times those in the control (Supplementary Table 1). On the other hand, we found 320 genes for which the mRNA levels in the mutant decreased to less than 1% of those in the control (Supplementary Table 2). Among the down-regulated genes, peak expression of most of them occurred during pupal and adult stages in males or in adult testes. We collected male larvae and prepared total RNAs from both controls and mutants, because the *mxc* gene is X-linked. These results are consistent with the previous finding that mutant larvae for the *mxc* gene lack germline cells (see Introduction). Among the up-regulated genes, 5% of the genes correspond to tumor-related genes such as *Pvf2* and *nimC1*. In addition, surprisingly, 24% plus 7% of the genes correspond to immunity-related genes involved in host defense against bacterial infection (Figure 2). In this study, we focused on the up-regulation of the immunity-related genes in the *mxc* mutant. We did not observe any microbial infection at detectable levels in the *mxc^mbn1^* larvae. These results led us to speculate that *Drosophila* innate immune pathways could be activated in response to tumor cells.

**Figure 2.**
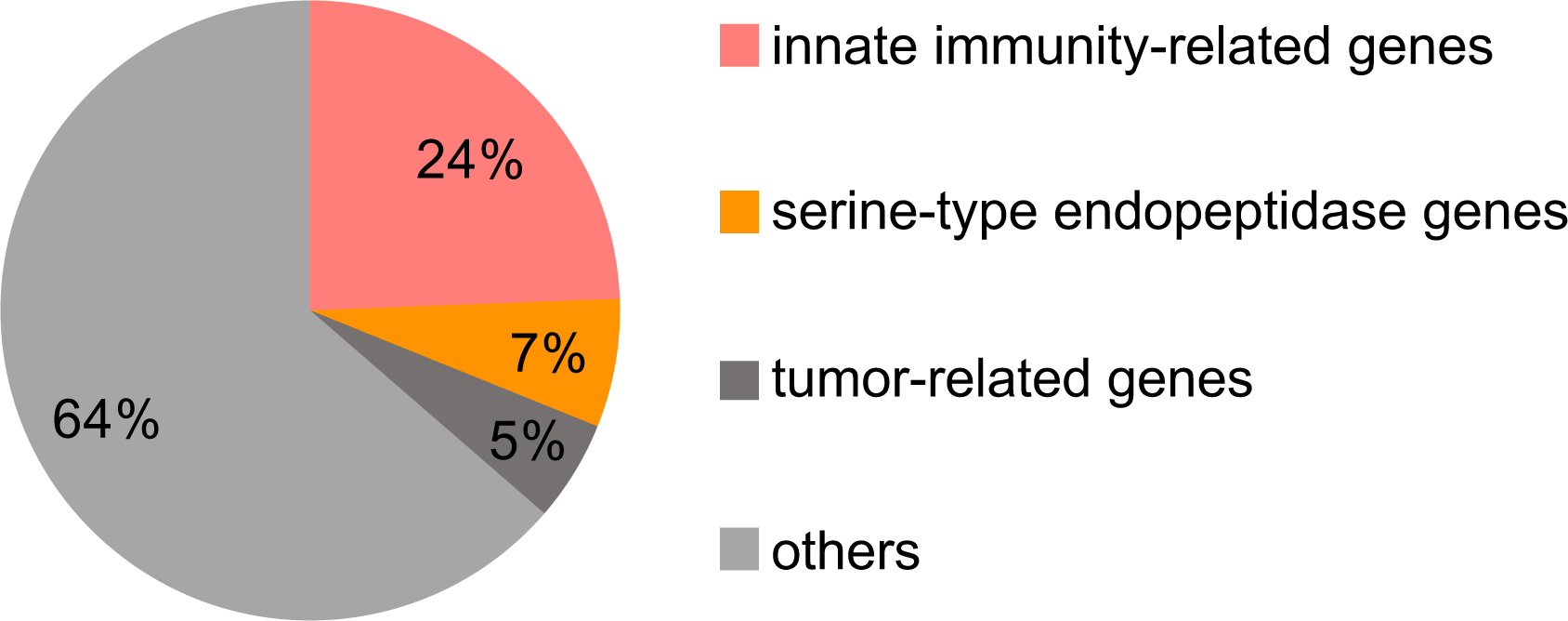
Identification of highly up-regulated genes in *mxc^mbn1^* mutants by RNA-Seq analysis. RNA-Seq analysis using total RNAs isolated from mature 3^rd^ instar larvae of wild-type males and *mxc^mbn1^* males. Among 209 genes whose mRNA levels increased more than 10-fold in the mutant larvae, immunity-related genes including endopeptidase genes account for 31% (24% plus 7%).

### Expression of target genes controlled by innate immune pathways is up-regulated in *mxc^mbn1^* mutant larvae

To confirm the RNA-Seq data and further investigate genes showing altered expression, we next tested whether three innate immune pathways, namely the Toll-mediated, Imd-mediated and JAK-STAT pathways, were activated in the mutant larvae harboring the LG tumors. First, we quantified relative mRNA levels of seven antimicrobial peptide genes, *Drs, Def, Diptericin* (*Dpt*), *Metchnikowin* (*Mek*), *Attacin A*(*AttA*), *Cecropin A2* (*CecA2*) and *Turandot B* (*TotB*), by qRT ‑ PCR (Figure 3A). Each of these genes is a target of one or two of the three innate immune pathways. We found that the amounts of these mRNAs increased in the *mxc^mbn1^* larvae by 6.9‒, 23.3‒, 3.4‒, 11.6‒, 5.0‒, 26‒ and 402-fold, respectively, compared with those in wild-type larvae (*p* < 0.0001). We failed to detect a similar up-regulation of any AMP gene in non-tumorous *mxc^G43^* mutant larvae, relative to wild-type whole larvae. Although a considerably higher expression of the *TotB* gene was seen even in *mxc^G43^* larvae, its mRNA level was only 4-fold higher in *mxc^mbn1^* larvae than in *mxc^G43^*.

**Figure 3.**
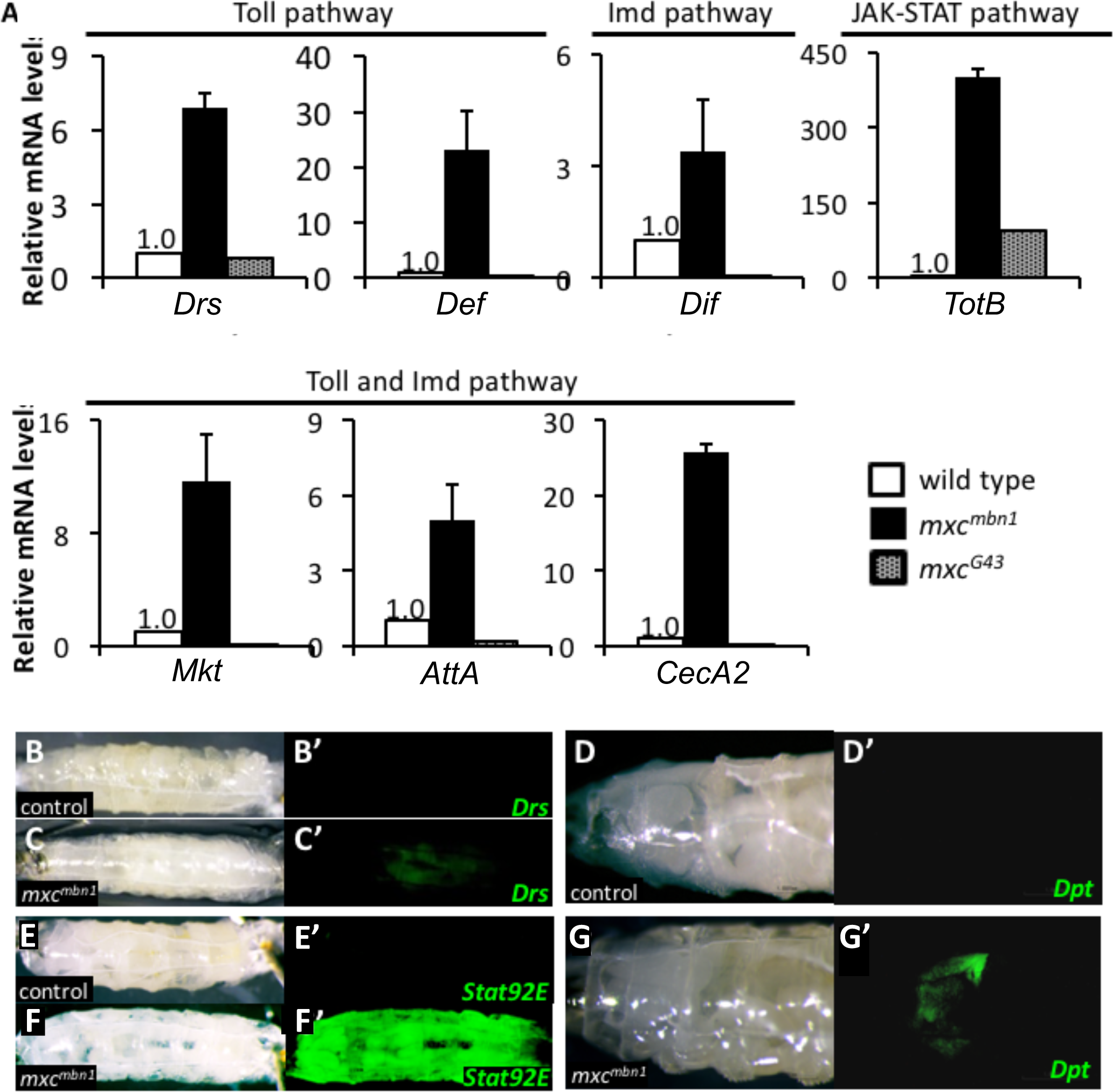
Activation of three innate immune pathways in the *mxc^mbn1^* mutant. (A) Relative mRNA levels of target genes of innate immune pathways in wild-type larvae, tumor-harboring larvae hemizygous for *mxc^mbn1^* and larvae hemizygous for non-tumorous *mxc^G43^*. mRNA was quantified by qRT-PCR. The mRNA levels for samples were normalized to wild-type values. Error bars represent SEM *Drs* and *Def* are controlled by the Toll pathway. *Dpt* is regulated by the Imd pathway. *Mtk*, *AttA* and *CecA2* are controlled by both Toll and Imd pathways. *TotB* is a target for the JAK-STAT pathway. Note that seven AMP genes are specifically up-regulated in the *mxc^mbn1^* mutant larvae. (B-G) Blight field images of control (B, D, E) and *mxc^mbn1^* (C, F, G) larvae. (B’, C’) Fluorescence images for *Drs-YFP* (Toll pathway) in control (B’) and *mxc^mbn1^* larvae (C’). Note that *Drs-YFP* was induced in fat bodies of the *mxc^mbn1^* mutant (C’), but not in control (B’). (D’, G’) Fluorescence images carrying *Dpt-YFP* (Imd pathway) in control (D’) and *mxc^mbn1^* larvae (G’). Note that *Dpt-YFP* was induced in fat bodies of the *mxc^mbn1^* mutant (G’), but not in control (D’). (E’, F’) Fluorescence images for *10×Stat92E-GFP* (JAK-STAT pathway) in control (E’) and *mxc^mbn1^* larvae (F’). Note that *10×Stat92E-GFP* was induced in fat bodies of the *mxc^mbn1^* mutant (F’), but not in control (E’).

Having obtained results indicating that expression of target genes activated by three immune pathways was significantly up-regulated in the mutant larvae, we further confirmed expression of the target gene products in the *mxc^mbn1^* larvae. Using three fluorescent protein reporters that monitor *Drs* (Figure 3B, C), *Dpt* (Figure 3D, G) and *Stat92E* gene-dependent transcription (Figure 3E, F), we were able to examine activation of the Toll-mediated, Imd-mediated and JAK-STAT pathways, respectively. In control larvae, expression of these targets was not stimulated in the absence of bacterial infection (Figure 3B’, 3D’, 3E’). On the other hand, expression of the *Drs* gene was substantially up-regulated in the fat body, which is a major tissue that produces and secretes AMPs, in *mxc^mbn1^* larvae (Figure 3C’). We also observed induction of *Dpt-YFP* in fat body as well as in other larval tissues in *mxc^mbn1^* (Figure 3G’). We detected GFP fluorescence generated from the *10×Stat92E-GFP* reporter (Figure 3F’), which monitors activation of the JAK/STAT signaling pathway in the mutant larvae. These results demonstrated that the Toll-mediated, Imd-mediated and JAK-STAT pathways were activated in the fat body of the *mxc^mbn1^* larvae without microbial infection.

### Ectopic activation of innate immune pathways suppresses the LG phenotype of *mxc^mbn1^*

The finding that *Drosophila* innate immune pathways were activated in the *mxc^mbn1^* larvae allowed us to test the possibility that activation of the innate immune pathways can influence tumor development. We examined whether up-regulation of these pathways could suppress the tumor overgrowth phenotype in *mxc^mbn1^* larvae. We induced ectopic overexpression of the Dl transcription factor acting in the Toll-mediated pathway, of a constitutively active form of Toll receptor, and of a ligand for the JAK-STAT signaling pathway. For ectopic induction of these factors in fat body, which is a major AMP-producing tissue, we used fat body-specific Gal4 drivers. With *ppl-gal4*, *r4-gal4* and *Lsp2-gal4,* which can induce *UAS*-dependent gene expression in fat body cells, the amounts of *Drs* mRNA increased by 26‒, 16‒ and 2.6-fold, respectively, compared with those in wild-type larvae. Consequently, we observed that ectopic activation of innate immune pathways suppressed the LG tumor overgrowth phenotype (Figure 4). The average size of the LG in *mxc^mbn1^/Y; Lsp2>Dl* was 34% of that in *mxc^mbn1^/Y*; *Lsp2>GFP* (n > 20), indicating that ectopic activation of the Toll pathway by *Dl* overexpression in fat body cells significantly suppressed the tumor overgrowth phenotype that appeared in *mxc^mbn1^* larvae (*p* < 0.0001, Student’s t test) (Figure 4C, 4F). Moreover, the average size of the LG of *mxc^mbn1^/*Y*; Lsp2>Toll^10B^* was 47% of that in *mxc^mbn1^/Y; Lsp2>GFP* (n > 20), indicating that ectopic activation of the Toll pathway by overexpression of the constitutively active form of a Toll receptor in fat body cells significantly suppressed the tumor overgrowth phenotype (*p* < 0.01, Student’s t test) (Figure 4B, D, F). Furthermore, the average size of the LG of *mxc^mbn1^/Y; ppl>Upd3* was 54% of that in *mxc^mbn1^/Y; ppl>GFP* (n>20), indicating that ectopic activation of the JAK-STAT pathway by *Upd3* overexpression in fat body cells significantly suppressed the tumor overgrowth phenotype (*p* < 0.001, Student’s t test) (Figure 4E and 4F). These observations indicate that ectopic activation of every innate immune pathway in the fat body can significantly suppress the LG tumor overgrowth phenotype in *mxc^mbn1^* larvae.

**Figure 4.**
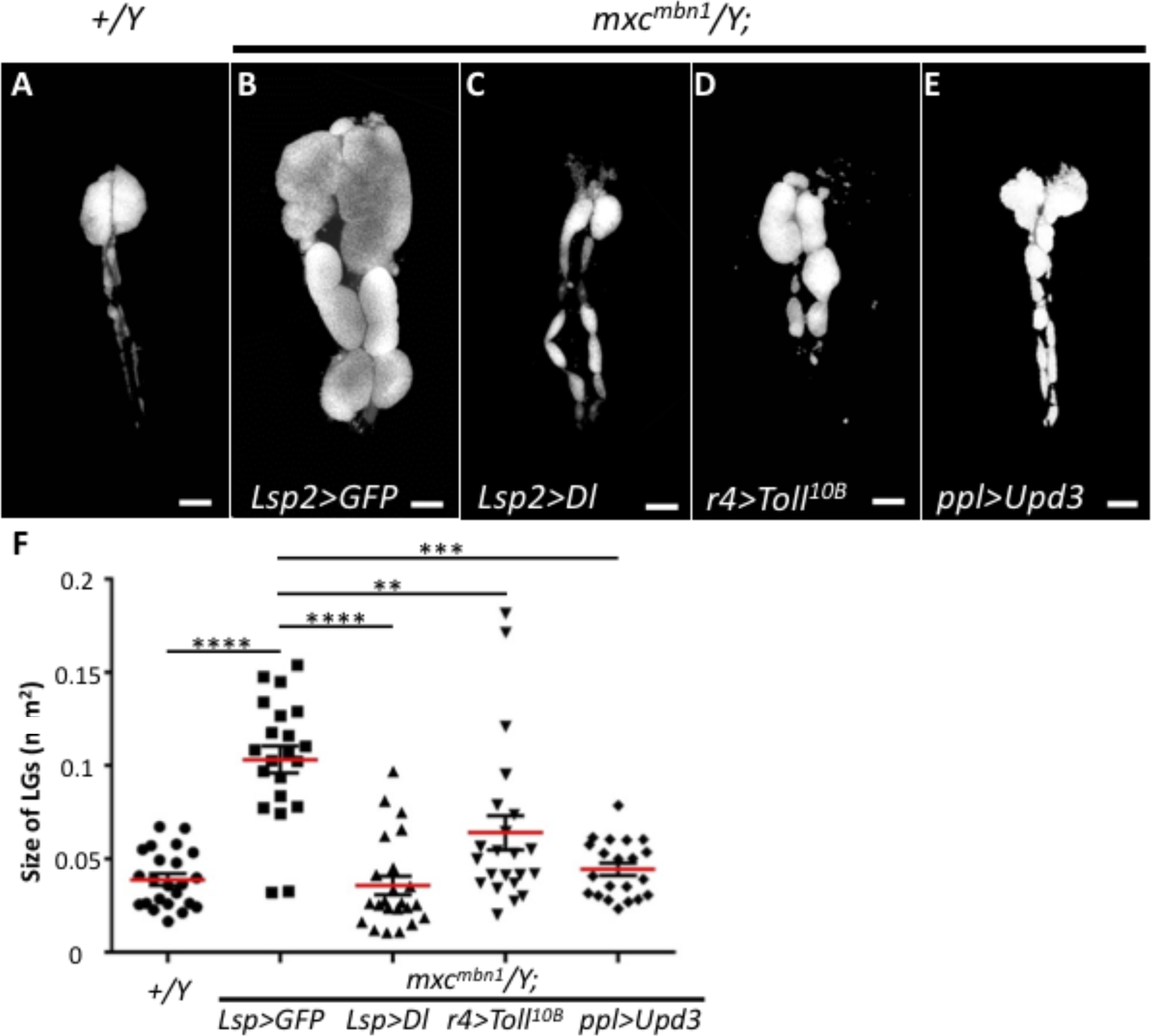
Overexpression of innate immune pathway components suppresses LG tumor overgrowth. (A–E) LGs from a wild-type larva (A), or from *mxc^mbn1^* larvae overexpressing GFP (B), *Dorsal* (C), *Toll^10B^* (constitutively active form of Toll) (D) and *Upd3* (E). (Bars: 100 μm.) (F) Quantification of tumor size (Student’s t test, \*\**p* < 0.01, \*\*\**p* < 0.001 and \*\*\*\**p* < 0.0001. Error bars represent SEM. Red horizontal lines represent average.).

Conversely, we tried to reduce the effect of each innate immune pathway by introducing mutations in genes required for these pathways. By genetic crosses, we generated *mxc^mbn1^* larvae carrying a heterozygous amorphic mutation for individual components of innate signaling pathways, namely *Dl*, *Toll*, *imd* or *Stat92E*, and then examined whether growth of the LG tumors changed under these conditions. We noticed that the reduced gene dose of the immune factors enhanced the LG tumor overgrowth phenotype in every case (Supplementary Figure 1A-F). The average size of LGs from *mxc^mbn1^/Y; dl^4^/+* larvae was about twice that of *mxc^mbn1^/Y* (n > 20) (*p* < 0.01, Student’s t test) (Supplementary Figure 1G). Also, LGs of *mxc^mbn1^/Y; Toll^rv1^/+* larvae were about twice as large as those of *mxc^mbn1^/Y* (n > 20) (*p* < 0.05, Student’s t test). The average size of LGs of *mxc^mbn1^/Y; imd/+* larvae was about four times that of *mxc^mbn1^/Y* (n > 20) (*p* < 0.0001, Student’s t test). The average size of LGs of *mxc^mbn1^/Y; Stat92E^HJ^/+* larvae was about twice that of *mxc^mbn1^/Y* (n > 18) (*p* < 0.05, Student’s t test). In summary, the LG tumor overgrowth phenotype was significantly enhanced in the *mxc^mbn1^* larvae carrying the mutations in factors of innate immune pathways. Taking these results together with the genetic data mentioned previously, we conclude that modification of the innate immune pathways, which can be activated in response to LG tumor cells, alters the tumor overgrowth phenotype of LG tumors in *mxc^mbn1^*.

### AMPs produced in the fat body suppress the LG tumor phenotype

The genetic data presented above allowed us to speculate that ectopic induction of the AMP genes, which are targets of the innate immune pathways, was responsible for the suppression of the LG tumor overgrowth phenotype in *mxc^mbn1^*. To test whether the AMPs have a suppressive effect on the LG tumor overgrowth, we induced ectopic expression of five different AMPs in fat bodies of *mxc^mbn1^* larvae using the fat body-specific *gal4* driver *r4-gal4*. This revealed a significant suppression of the LG tumor overgrowth phenotype in *mxc^mbn1^* by expression of every AMP (Figure 5B-G, 5H). The average size of the LGs in *mxc^mbn1^/Y; r4>Drs* was 58% of that in *mxc^mbn1^/Y; r4>GFP* (n > 55), indicating that ectopic *Drs* overexpression in fat body cells significantly suppressed the tumor overgrowth phenotype (*p* < 0.0001, Student’s t test) (Figure 5B and 5C). Moreover, the average LG size in *mxc^mbn1^* mutants similarly expressing the other four AMPs decreased to 48% to 70% of the average size in *mxc^mbn1^* mutants without AMP overexpression (n>30, *p* < 0.01, Student’s t test) (Figure 5H). These observations strongly suggest that suppression of LG tumor overgrowth in *mxc^mbn1^* by activation of the three innate immune pathways is a consequence of ectopic expression of AMPs in fat bodies. Considering the genetic results described here, we conclude that the *Drosophila* innate immune pathways can be activated in response to the LG tumor cells and that the expression of their target genes encoding AMPs can suppress the LG tumor growth.

**Figure 5.**
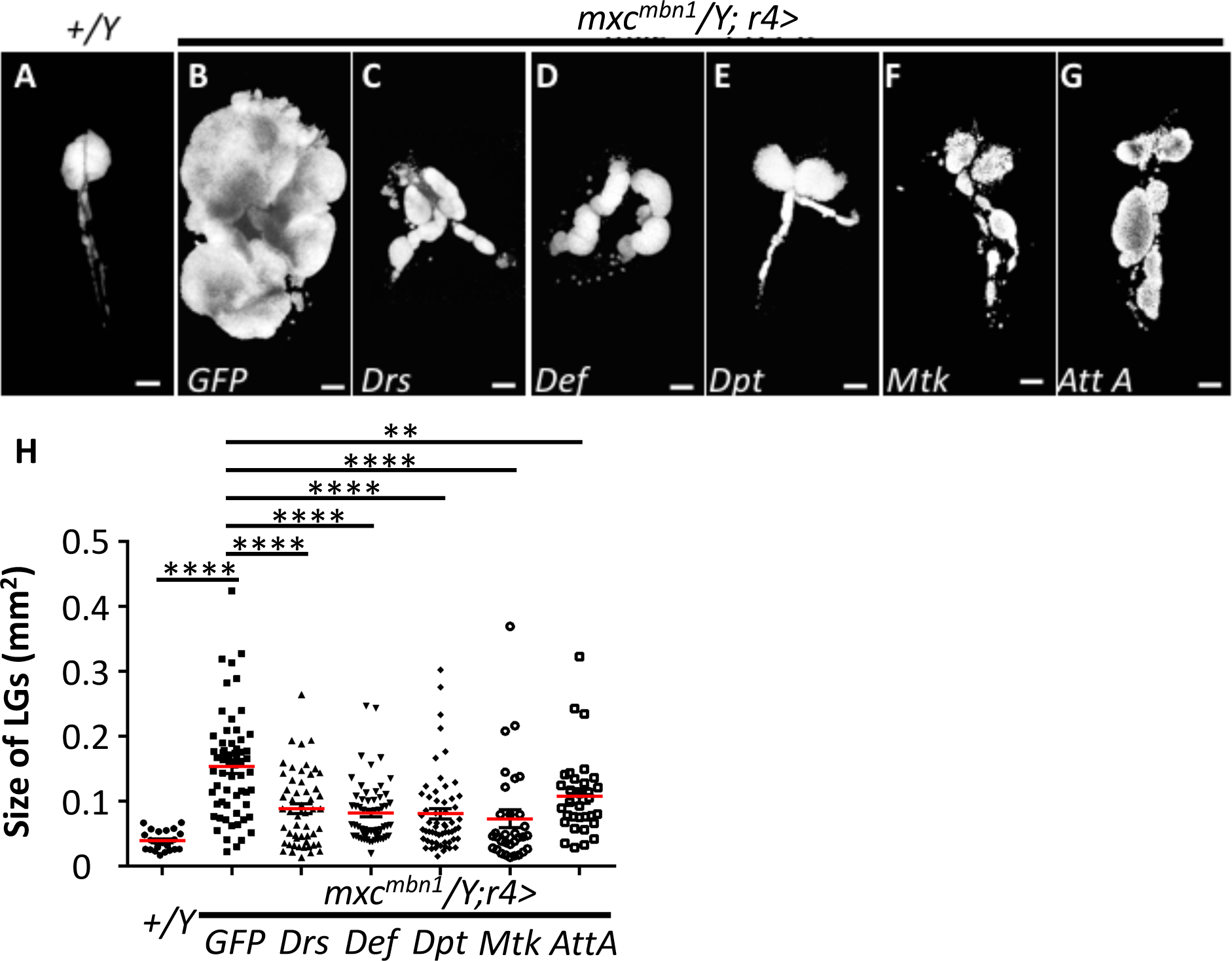
Fat body-specific induction of AMP genes that are targets of innate immunity pathways in *mxc^mbn1^* larvae. (A–G) LGs from a wild-type larva (A), or from *mxc^mbn1^*mutant larvae expressing GFP (*mxc^mbn1^; r4>GFP*) (B), *Drs* (*mxc^mbn1^; r4>Drs*) (C), *Def* (*mxc^mbn1^; r4>Def*) (D), *Dpt* (*mxc^mbn1^; r4>Dpt*) (E), *Mtk* (*mxc^mbn1^; r4>Mtk*) (F) and *AttA* (*mxc^mbn1^; r4>AttA*) (G). (Bars: 100 μm.) (H) Quantification of tumor size (Student’s t test, \*\**p* < 0.01 and \*\*\*\**p*

### Ectopic induction of AMPs in the fat body enhances apoptosis in the *mxc^mbn1^* LG, but not in normal tissues

To test the hypothesis that AMPs encoded by target genes of innate immune pathways are responsible for the suppressive effect on development of the LG tumors in *mxc^mbn1^* larvae, we examined whether AMPs could stimulate apoptosis in the tumor cells. LG cells undergoing apoptosis in mature larvae were detected using the Cell Event Caspase-3/7 Green Detection assay. No apoptotic signals were seen in normal LGs in control larvae expressing the *Drs* gene in fat body (n = 24) (Figure 6A, A’). In contrast, we observed apoptosis signals in 7.7% of the whole LG region in *mxc^mbn1^* larvae at the wandering stage of 3^rd^ instar (n = 20) (Figure 6B, B’, F). Interestingly, in *mxc^mbn1^* larvae at the same larval stage over expressing *Drs*, *Def* and *Dpt* specifically in fat bodies, more apoptosis signals were observed over a wider region of the whole LG (14.8%, n = 20 (*mxc^mbn1^/Y; r4>Drs*), 17.8%, n = 18 (*mxc^mbn1^/Y; r4>Def*), 16.3%, n = 22 (*mxc^mbn1^/Y; r4>Dpt*) (Figure 6C-E, 6C’-E’, 6F), compared with *mxc^mbn1^* without induced AMP expression. However, we never saw any apoptotic signals in other larval tissues in the mutants, including the larval central nervous systems, wing, haltere and eye discs. Moreover, no apoptosis signals appeared in control larvae overexpressing *Drs* or *Dpt* in their fat bodies (n > 9) (Figure 6A). However, the overexpression of every AMP failed to rescue the larval mortality in *mxc^mbn1^* larvae (n <100 larvae), since it had no effects on the reduced tissue growth in their imaginal discs (Figure 6G-L). From these results, we concluded that the AMPs had a cytotoxic effect that induced apoptosis exclusively in the tumor cells.

**Figure 6.**
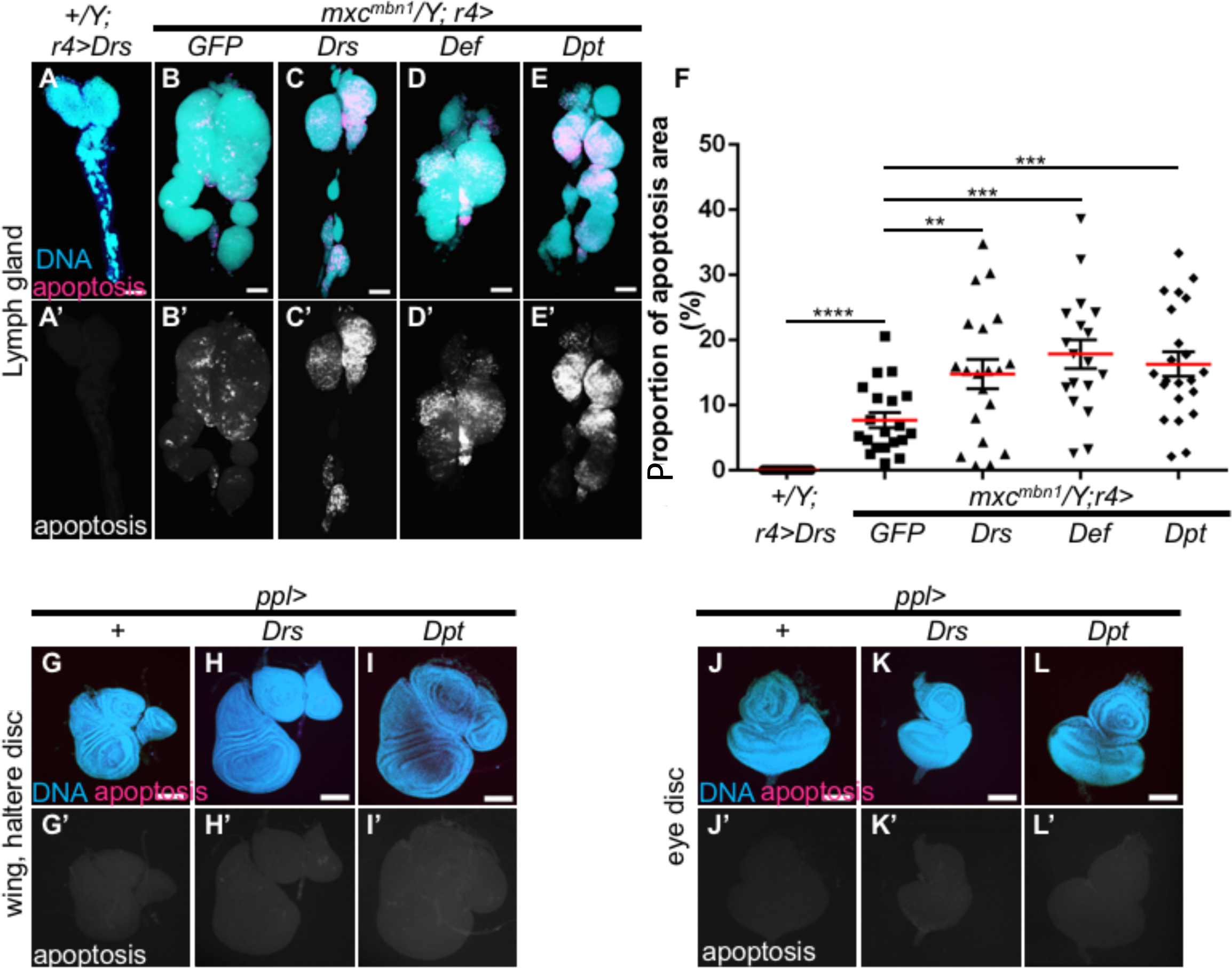
Observation of LG cells undergoing apoptosis in *mxc^mbn1^* larvae overexpressing AMPs. (A-E, G-L) Cell Event Caspase-3/7 Green Detection Reagent assays in LGs, wing, haltere discs and eye discs. Red signals in A-E and G-L, and white signals in A’-E’ and G’-L’, represent apoptotic cells. Blue, DNA. (A-E) Immunofluorescence of LG. (A, A’) Control LG expressing *Drs* (*r4>Drs*). Note that no apoptotic cells are evident. LGs of *mxc^mbn1^* larvae expressing GFP (B, B’), *Drs* (C, C’), *Def* (D, D’) and *Dpt* (E, E’) in their fat bodies. The LG of *mxc^mbn1^* contains many apoptosis signals (B, B’). LGs of *mxc^mbn1^* larvae overexpressing *Drs* (C, C’)*, Def* (D, D’) and *Dpt* (E, E’) show more apoptosis signals than *mxc^mbn1^* larvae. Bars: 100 μm. (F) Proportion of apoptosis area in total LG area (Student’s t test, \*\**p* < 0.01, \*\*\**p* < 0.001, \*\*\*\**p* < 0.0001. Error bars represent SEM. Red horizontal lines represent average.). (G-I) Immunofluorescence of wing and haltere discs. (J-L) Immunofluorescence of eye discs. No apoptosis signals can be seen in wing or haltere and eye discs in control larvae overexpressing *Drs* or *Dpt* genes in their fat bodies.

### Preferential association of AMPs with regions showing a loose cell-to-cell adhesion and possessing lower levels of DE-cadherin in the LG tumors

As AMPs secreted into the hemolymph can induce apoptosis in the LG tumor cells but not in normal LG cells, we next addressed the mechanism underlying the observation that apoptosis induction specifically occurred in the tumor LGs. AMPs are mainly synthesized in the fat body cells and secreted into the hemolymph. Therefore, these AMPs act on the LG tumors via the hemolymph in *mxc^mbn1^* larvae. We induced expression of Drosomycin, Defensin and Diptericin fused with a HA-tag in fat bodies in control *mxc^mbn1^* mutant larvae (*r4>AMP-HA* and *mxc^mbn1^/Y; r4>AMP-HA*). These tagged AMPs produced in and secreted from the fat bodies were detected by anti-HA antibody immunostaining. We failed to find any signals in control LGs (*r4> Drs-HA*, *r4>Def*-HA, *r4>Dpt-HA*) (Figure 7A, B, E, F, I), except in pericardial cells, which play a role in pumping hemolymph in the open circulatory system (asterisks in Figure 7A, C, E, G, I, K). The presence of Drosomycin in pericardial cells suggests that these AMPs in hemolymph are preferentially incorporated into the pericardial cells. On the other hand, we saw distinct immunostaining signals of these three AMPs in the LG tumors in *mxc^mbn1^* larvae overexpressing the AMPs with a HA-tag (Figure 7C, G, K). In particular, we observed the AMP signals along LG regions in the mutant, which showed reduced cell density (Figure 7C’, G’, K’) and a reduced DE-cadherin distribution in *mxc^mbn1^* (Figure 7C’’, G’’, K’’). Finer observation of the LG samples at high magnification suggested that Drosomycin and Defensin were taken up by hemocyte-like cells associated with the LG regions in *mxc^mbn1^* (Figure 7D, H). Many small punctate signals of these two AMPs were also observed in the LGs. Diptericin was similarly associated with the LG, forming fibrous short fragments (Figure 7L). Even in controls overexpressing *Dpt*, similar short fragments as well as many small dots containing the AMP were found in dorsal vessels, but not in the LGs (Figure 7J).

**Figure 7.**
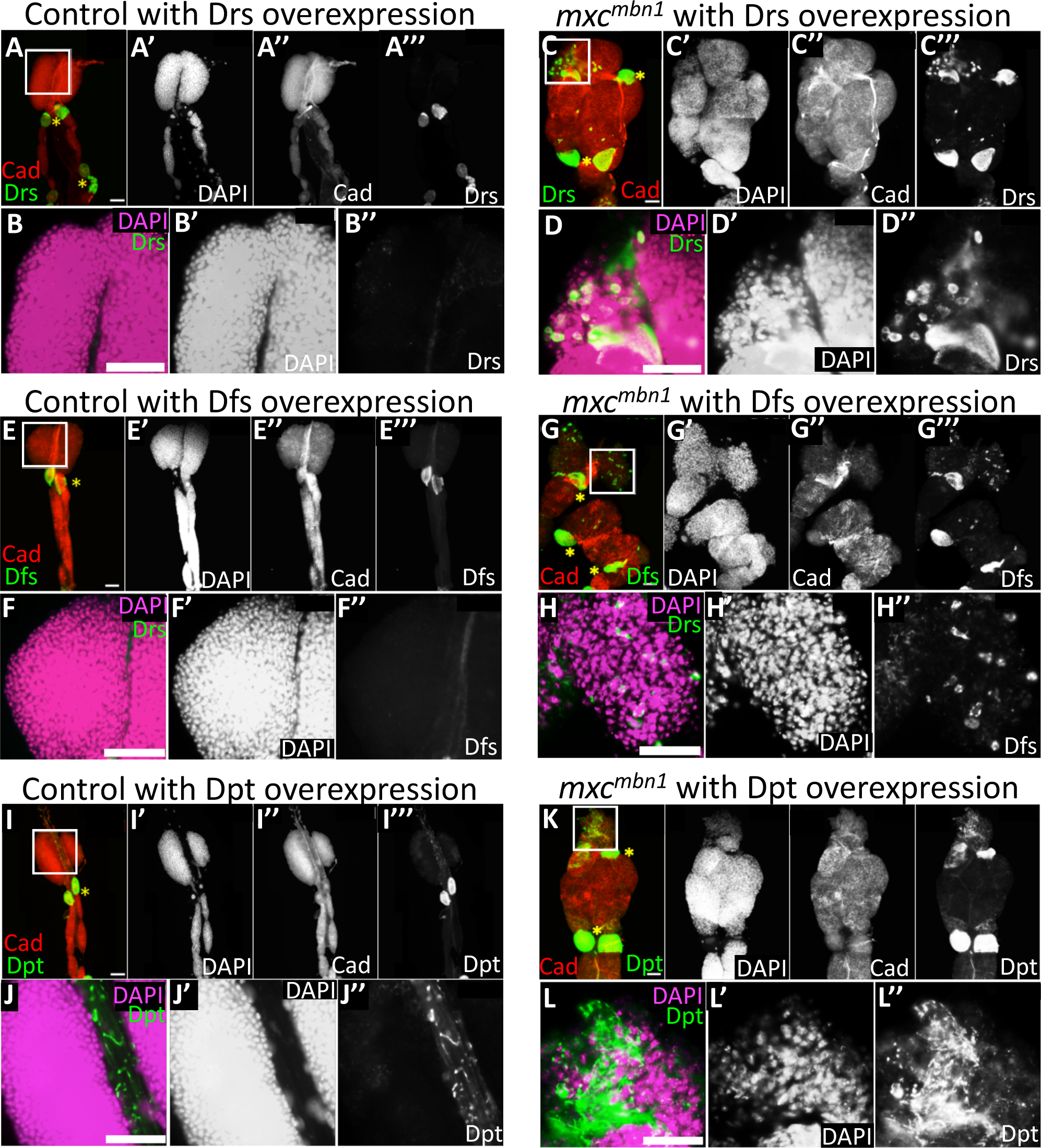
Visualization of AMP on LGs and in circulating hemocytes from normal control and *mxc^mbn1^* larvae overexpressing AMPs. (A–H) Localization on LGs of AMPs fused with HA-tag. (A, B) Control LG (*r4>Drs*), (C, D) LGs from *mxc^mbn1^* overexpressing *Drs* (*mxc^mbn1^; r4>Drs*). (E, F) Control LG (*r4>Def*), (G, H) LGs from *mxc^mbn1^* overexpressing *Def* (*mxc^mbn1^; r4>Def*). (I, J) Control LG (*r4>Dpt*), (K, L) LGs from *mxc^mbn1^* overexpressing *Dpt* (*mxc^mbn1^; r4>Dpt*). (A, C, E, G, I, K) Red: DE-cadherin (Cad); green: AMP (drosomycin, defensin or diptericin). (B, D, F, H, J, L) High-magnification views. Red: DNA; green: AMP. ⋇ Pericardial cells. AMPs are often localized on regions exhibiting reduced DE-cadherin immunostaining signals. Drosomycin and Defensin appear to be taken into hemocyte-like cells (D, H), whereas diptericin is associated with the LG in a fibrous shape (H). (Bars: 50 μm.)

Because we found hemocyte-like cells containing Drosomycin and Defensin on the LG tumors in *mxc^mbn1^* larvae, we next examined whether circulating hemocytes in the hemolymph also contained AMPs. We induced three AMPs exclusively in larval fat bodies and examined their accumulation in circulating hemocytes by anti-HA-tag immunostaining. More intense anti-HA immunostaining signals appeared in the cytoplasm of circulating hemocytes in tumorous *mxc* mutant (*mxc^mbn1^/Y; r4>AMP-HA*) (Figure 8B, D, F) than in controls (*r4>AMP-HA*) (Figure 8A, C, E). *Def* mRNAs increased 753-fold (*r4>Def*) and 716-fold (*mxc^mbn1^/Y; r4>Def*) under the *r4-Gal4* driver, compared with the level in wild-type larvae (Supplementary Figure 2A). These data indicate almost the same expression levels of *Def* in fat bodies. Nevertheless, hemocytes in *mxc^mbn1^* larvae contained much higher amounts of AMPs. Similarly, *Dpt* mRNAs were also up-regulated 140-fold (*r4>Dpt*) and 226-fold (*mxc^mbn1^/Y; r4>Dpt*), relative to wild-type larvae (Supplementary Figure 2B). Although the mRNA level of *Dpt* in *mxc^mbn1^* larvae was 60% higher than that in controls, circulating hemocytes in *mxc^mbn1^* displayed more intense fluorescence signals of the AMP. Taken together, these results led us to the following interpretation: AMPs produced by the fat body are secreted into the hemolymph in response to LG tumors in *mxc^mbn1^* larvae. Subsequently, the AMPs in hemolymph are incorporated into circulating hemocytes and become preferentially associated with regions showing reduced cell-cell adhesion.

**Figure 8.**
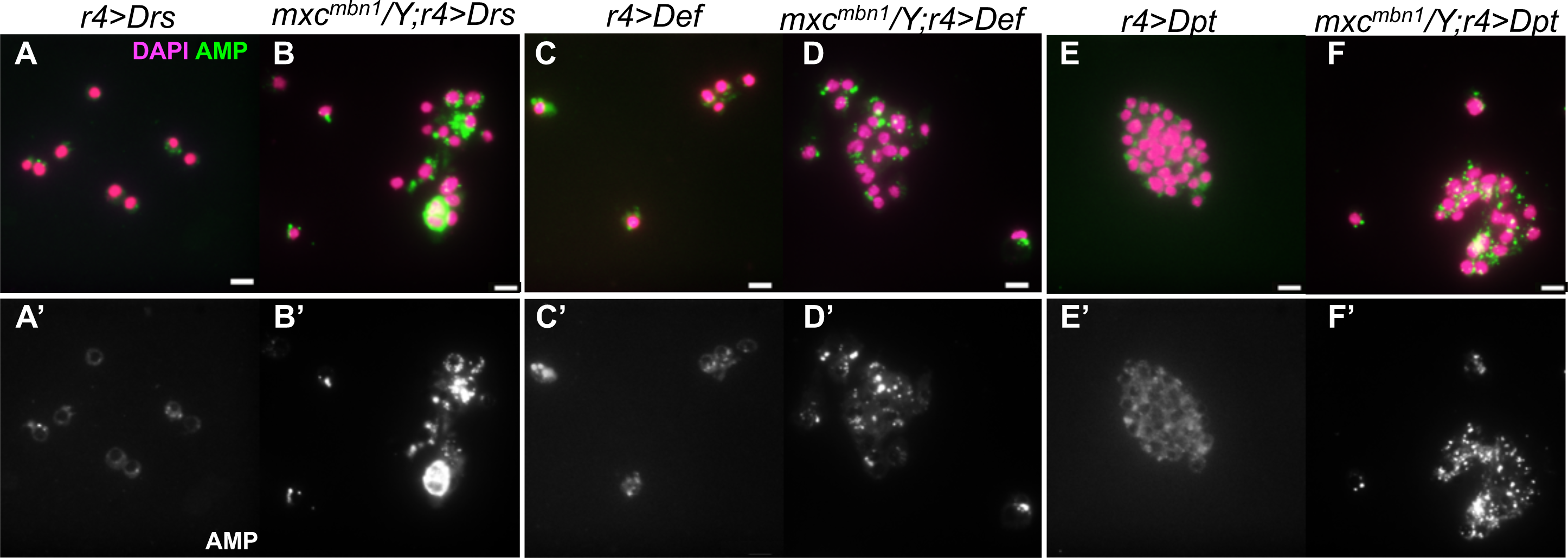
Visualization of AMP on LGs and in circulating hemocytes from normal control and *mxc^mbn1^* larvae overexpressing AMPs. (A-F) Localization of AMPs fused with HA-tag in circulating hemocytes. (A) Circulating hemocytes from control (*r4>Drs*) and (B) *mxc^mbn1^; r4>Drs*. (C) Circulating hemocytes from control larvae (*r4>Def*) and (D) mutant larvae (*mxc^mbn1^; r4>Def*). (E) Circulating hemocytes from control larvae (*r4>Dpt*) and (F) mutant larvae (*mxc^mbn1^; r4>Dpt*). Red: DNA; green: AMP. (Bars: 10 μm.)

## Discussion

In this study, we have examined whether the innate immune system can suppress a malignant blood neoplasm phenotype observed in a larval hematopoietic tissue termed the lymph gland (LG) of *Drosophila mxc^mbn1^* mutants. To identify genes whose mRNA levels were altered in larvae having the LG tumor, we first carried out a comprehensive RNA-Seq analysis. This indicated that many immune response genes were up-regulated in the mutant larvae. *Drosophila* has three immune pathways, namely the Toll-mediated, Imd-mediated and JAK-STAT pathways. Upon activation of these pathways, expression of their target genes, *Drs*, *Def*, *Dpt*, *Mtk*, *AttA*, *CecA2 and TotB*, encoding antimicrobial peptides is induced. By qRT-PCR analysis, we demonstrated that the mRNA level of all seven AMP genes was significantly increased. Using GFP-reporters to visualize gene expression, we further confirmed that expression of the target genes was induced in the mutant fat bodies, which are responsible for AMP production. The LG tumor phenotype in the *mxc* mutants was significantly enhanced in *mxc^mbn1^* larvae carrying mutations in the *Toll* and *imd* genes. Furthermore, we demonstrated that ectopic expression of any of five AMPs in fat bodies can significantly suppress the tumor phenotype, although the ectopic expression failed to rescue the larval lethality in the mutants. Simultaneously, it stimulated apoptosis in the LG tumors in *mxc^mbn1^*, whereas no apoptosis occurred in control LGs. Furthermore, we addressed the mechanism underlying apoptosis induction specifically in the tumor LGs. We found a distinct localization of the AMPs on the LG tumors in *mxc^mbn1^*, but not on normal LGs. Both Drosomycin and Defensin appeared to be taken up in hemocyte-like cells associated with the LG regions in *mxc^mbn1^*, whereas Diptericin was associated with the LG without intervening hemocytes, inducing the formation of fibrous short fragments. These results suggest that the AMPs can associate with LG tumors in different ways, being either incorporated into circulating hemocytes or not.

### Innate immune systems can suppress the development of tumors generated in a larval hematopoietic tissue in *Drosophila*

Expression of all genes encoding antimicrobial peptides rose simultaneously in the tumorous *mxc^mbn1^* mutants. These molecular data indicate that all three innate immune pathways in *Drosophila* were activated in *mxc^mbn1^* mutants carrying the LG tumors. This activation was required for suppression of tumor growth. Moreover, we obtained direct evidence that ectopic expression of AMPs in fat bodies significantly suppressed the tumor phenotype. These genetic data led us to hypothesize that there exists a regulatory mechanism by which the innate immune signaling pathways are activated in response to hematopoietic tumors and induce expression of the AMP genes. It thus seems reasonable to speculate that the AMPs have anti-cancer effects that can suppress hyper-proliferation of tumor cells. Our genetic data imply that a defense mechanism against tumor cells exists even in insects, despite their lacking an acquired immune system. If the innate immune system in insects plays a role as a primitive prophylactic against tumor cells, it would be advantageous to organisms lacking the more advanced immune system that has been acquired during evolution by vertebrates (Hoffmann and Reichhart, 2002). Because tumorigenesis is probably an ineluctable feature for multicellular organisms (Domazet-Lošo et al., 2014), it is important to suppress the unchecked growth of abnormal cells that deviate from their proper role in a multicellular system. However, few biological studies that address the issue of how invertebrates challenge tumor cells have been conducted. Interestingly, our finding that AMPs in the hemolymph enhanced apoptosis exclusively on the tumors raises the possibility that *Drosophila* has a biological defense system that can distinguish tumor cells from normal cells. A recent study has also shown that epithelial solid tumors induce activation of Toll signaling in fat body cells, which causes cell death of epithelial tumors and a reduction in their size (Parisi et al., 2014). However, mechanism(s) underlying a suppression of tumor growth in invertebrates have hardly been investigated to date. In this study, we have for the first time uncovered genetic evidence that AMPs play a crucial role in anti-tumor activity of the innate immune system against tumor development in *Drosophila*. Since AMPs exist in many and diverse species, it is reasonable to anticipate that our current findings in *Drosophila* will be pertinent to numerous other organisms. While the acquired immune system in mammals is more effective for preventing cancer progression, it is also possible that their innate immune system makes a greater contribution than is usually assumed. For example, it may play a major role in an initial attack against transformed cells (Kim et al., 2007). If this is the case, the AMPs may lead to a breakthrough for basic research on cancer immunotherapy in the near future.

### Drosomycin and Defensin secreted from fat bodies into the hemolymph act on hematopoietic tumors via circulating hemocytes

Ectopic expression of five AMPs in fat bodies of the *mxc^mbn1^* mutants significantly suppressed the LG tumor phenotype, and also enhanced apoptosis in the tumorous LG of *mxc^mbn1^*. We saw abundant AMPs in these tissues, but not in the controls. Interestingly, Drosomycin and Defensin appeared to enter hemocyte-like cells that were associated with LG regions showing a more severe tumor phenotype. In addition, much higher AMP levels were found in the cytoplasm of circulating hemocytes in the *mxc* mutant larvae. These observations imply that Drosomycin and Defensin were preferentially taken into circulating hemocytes in the *mxc* mutant larvae, compared with the controls. This is consistent with a previous observation that some AMPs have concentration-dependent cytotoxic effects on mammalian tumor cells and normal cultured cells (Hoskin and Ramamoorthy, 2008). AMPs produced in the fat body are secreted into the hemolymph (Lemaitre and Hoffmann, 2007). Surprisingly, we observed that circulating hemocytes containing AMPs were associated with the LG tumors. However, previous studies have reported that *Drosophila* AMPs stored in the cytoplasmic granules of circulating hemocytes were discharged against infecting microbes (Destoumieux et al., 2000; Yakovlev et al., 2017). Moreover, AMP-containing hemocytes are reportedly recruited to neoplastic tumors in *Drosophila* (Cordero et al., 2010; Parisi et al., 2014; Pastor-Pareja et al., 2008). Subsequently, the hemocytes release the stored AMPs against the tumor cells by regulated exocytosis (Destoumieux et al., 2000; Yakovlev et al., 2017). In this study, we showed that Drosomycin and Defensin have an anti-tumor activity against hematopoietic tumors with the help of circulating hemocytes in the hemolymph. From these findings, another new issue arises: can the AMPs influence tumor growth in tissues other than the lymph gland? In our current system, it is difficult to investigate interactions between the hemocytes and cells in the tumorous tissues because both cell types share a common origin in development. Therefore, it would be interesting to examine the effects of AMPs against tumors generated in different tissues, such as imaginal discs.

### How does Diptericin specifically target tumor cells?

In contrast to the mechanism of action of Drosomycin and Defensin against the LG tumors as described above, Diptericin seems to exert its anti-tumor activity by a different mechanism: it apparently acts more directly on the tumors, without the intervention of circulating hemocytes. The effect of Diptericin is restricted to the tumor cells and does not extend to normal cells. A hypothesis has been proposed in which AMPs have a higher affinity against tumor cells than against normal cells because of a difference in electric charge on the plasma membrane (Rivero-Muller et al., 2007). Some tumor cells have a net negative charge on the plasma membrane, similar to bacterial cells. The membranes of these tumor cells contain a higher concentration of anionic molecules such as phosphatidylserine and O-glycosylated mucins (Hoskin and Ramamoorthy, 2008; Mader and Hoskin, 2006; Papo and Shai, 2005). To examine whether the *mxc^mbn1^* LG tumors showed this characteristic, we carried out staining experiments using the fluorescent dye PSVue550 as a probe known to bind to several negatively charged phospholipids (Gray, 2011). However, we failed to detect any difference in staining intensity between the tumor LGs from *mxc^mbn1^*and normal LGs (data not shown). Therefore, it is necessary to consider a new mechanism for the restricted association of diptericin with LG tumors in future studies.

On the basis of the apoptosis assay, we concluded that AMP treatment enhanced apoptosis in the LG tumors. However, a mechanism for this induction has not yet been established. It has been argued that AMPs bind to tumor cell membranes via an electrostatic interaction and align themselves parallel to the membrane surface (Rivero-Muller et al., 2007). After enough peptide molecules have assembled on the membrane surface, they permeate the membrane, which is followed by membrane disintegration and micellization. Finally, pores spanning the membrane are transiently formed, and this leads to cell death. Once AMPs are taken into the cell, they can perturb the integrity of mitochondrial membranes having a negative charge, resulting in the release of various mitochondrial proteins including cytochrome *c*, a potent apoptosis inducer (Hoskin and Ramamoorthy, 2008; Mader and Hoskin, 2006). It is well known that release of cytochrome *c* from damaged mitochondria induces Apaf-1 oligomerization, caspase 9 activation and subsequently the conversion of pro-caspase 3 to caspase 3, which is responsible for many of the hallmarks of apoptotic symptoms (Porter and Jänicke, 1999; Zou et al., 1999). The mechanism underlying the association between AMP overexpression and apoptosis induction should be elucidated in further studies.

### Concluding remarks

In this study, we demonstrated that AMPs were induced as a consequence of activation of the innate immune pathways in response to hematopoietic tumors in *Drosophila*. We obtained genetic evidence that individual AMPs enhanced apoptosis in the tumors by two different routes, either with the aid of circulating hemocytes or by direct binding to the tumors. As a variety of AMPs are found in many organisms from plants to human (Hoskin and Ramamoorthy, 2008; Mader and Hoskin, 2006), they are regarded as common protection factors that are essential for an initial attack against a microbial infection by the innate immune system. They are involved in the immune response not only against microbial pathogens but also against tumor cells in both mammals and *Drosophila* (Winder et al., 1998; Zaiou, 2007; this study). Some human AMPs also display anti-cancer activity that induces cell death in cancer cells (Baker et al., 1993; Sloballe et al., 1995). Considering these features, AMPs are promising as new anti-cancer drugs. Our findings that some AMPs act on tumor cells through circulating hemocytes, and while another assembles to form unique structures and thus acts directly on tumor cells, offer an insight into understanding the mechanism of their specific effects on tumor cells. Besides selective cytotoxicity, the deployment of AMPs has many potential advantages in cancer chemotherapy. It is unlikely that AMPs bind to specific extracellular or intracellular receptors. Therefore, they should represent superior anti-cancer drugs, because cells acquiring resistance to such agents rarely appear (Gaspar et al., 2013). One of the most serious problems in cancer chemotherapy is the strong side-effects of anti-cancer drugs. In this context, our data indicate that overexpression of three AMPs had no adverse effects in *Drosophila*. Considering previous and present results, the AMPs are attractive candidates as therapeutic agents for cancer treatment. However, because most biological studies on the AMPs have been performed using cultured cell lines, it is now essential to obtain data on AMPs at the organism level. The experimental system established in this study allows us to investigate the anti-cancer effects of AMPs in *Drosophila*, and should therefore become a fascinating model in future cancer research.

## Materials and Methods

### *Drosophila* stocks

All *Drosophila melanogaster* stocks were maintained on standard cornmeal food, as previously described (Oka et al., 2015). Canton S was used as a wild type control stock. A recessive lethal allele of *mxc* showing the LG tumor phenotype, *mxc^mbn1^* was obtained from Bloomington Drosophila Stock Center (Bloomington, IN, USA). A hypomorphic allele, *mxc^G43^* not displaying any tumor phenotype were obtained from Kyoto Stock Center (Kyoto, Japan). To induce expression of AMP genes, Drs, Def, Dpt by Gal4/UAS system, we used the following the UAS stocks; *M{UAS-Drs.ORF.3xHA.GW}* (&F002219), *M{UAS-Def.ORF.3xHA.GW}* (&F002467) and *M{UAS-Dpt.ORF.3xHA.GW}* (&F002330) from the FlyORF (Zurich, Switzerland). Other UAS-AMP stocks for induction of Metchnikowin and Attacin A were gifts from Dr. B. Lemaitre (Swiss Federal Inst. Tech.). For genetic analysis to modify activity of innate immunity pathways, we used the following mutants, *dl^4^* (&BL7096; Bloomington Drosophila Stock Center), *Toll^1-rxa^* (a gift from Dr. Y. Yagi, Nagoya Univ.), *imd^1^* (a gift from Dr. Y. Yagi) and *Stat92E^HJ^* (&BL24510; Bloomington Drosophila Stock Center). For overexpression of innate immunity genes, we used *UAS-dl-3HA* (&F000638; FlyORF), *UAS-Toll^10B^* (&BL58987; Bloomington Drosophila Stock Center) and *UAS-Upd3* (a gift from Dr. B. Lemaitre). The following Gal4 driver stocks were used for ectopic expression in specific larval tissues; *ppl-Gal4* for expression in the fat body as well as in lymph gland at a higher level (a gift from Dr. P. Léopold, Institute Valrose Bidogie), *P{w[+mC]=r4-GAL4}3* (*r4-Gal4*) for fat body-specific expression at a moderate level (&BL33832; Bloomington Drosophila Stock Center), *P{Lsp2-GAL4}3* (*Lsp2-Gal4*) for fat body-specific expression in the 3rd instar larvae at a weaker level (&BL6357; Bloomington Drosophila Stock Center). *He-Gal4* for expression restricted to subpopulations (70-80 %) of the circulating hemocytes (a gift from Dr. D. Hultmark, Umea Univ.). To monitor activation of innate immune pathways, we used the following GFP reporter for AMP genes, *Drosomycin-YFP* (*Drs-YFP*) (a gift from Dr. Y. Yagi), *Diptericin-YFP* (*Dpt-YFP*) (a gift from Dr. Y. Yagi) and *PBac{y+mDint2 w+mC=Stat92E-GFP.FLAG}* (*Stat92E-GFP*) (&BL38670, Bloomington Drosophila Stock Center).

### Lymph gland (LG) preparation

To compare the size of LGs, we isolated a whole region of the LG from mature stage larvae and fixed them with 3.7 % folmaldehyde for five minutes. The fixed samples were mildly squashed using an apparatus so that the tissue spreaded into a few cell layers with a constant thickness.

### RNA-sequence analysis

Total RNA was extracted from 3rd instar larvae using the Trizol^®^ reagent (Invitrogen, Carlsbad, CA, USA). The isolated RNAs were used for the construction of single-end mRNA-seq libraries, using a NEBNext Ultra Directional RNA Library Prep Kit (New England Biolabs, Ipswich, MS, USA) according to the manufacturer’s recommendations. mRNA-seq was performed on an Illumina Hi-seq 1000 instrument (Illumina, San Diego, CA, USA) using 51-bp single-end reads. Read quality was checked for each sample using FASTQC (http://www.bioinformatics.babraham.ac.uk/projects/fastqc) (version 0.11.5). Filterd reads were aligned to the reference *D. melanogaster* genome sequence (BDGP5) using the TopHat program version 2.0.6 with default parameters (Kim et al., 2013). Cufflinks (version 2.0.2) was employed with default parameters for transcript assembly (Trapnell et al., 2012). The expression level of each gene was quantified as FPKMs (fragments per kilo base of exon per million mapped fragments). We carried out gene ontology (GO) analysis of genes expressed in adults and aligned the genes whose expression was significantly changed, with q-values of < 0.01. The GO classification system was applied by employing the database for annotation, visualization and integrated discovery (DAVID) version 6.7 (http://david.abcc.ncifcrf.gov/).

### qRT-PCR analysis

Total RNA was extracted from 3rd instar larvae with each genotype using the Trizol reagent (Invitrogen). cDNA synthesis from the total RNA was carried out using the PrimeScriptTM High Fidelity RT-PCR Kit (TaKaRa, Clontech Laboratories, Shiga, Japan) using an oligo dT primer. Real-time PCR was performed using the FastStart Essential DNA Green Master (Roche, Mannheim, Germany) and a Light Cycler Nano instrument (Roche). qPCR primers were synthesized as follows: RP49-Fw, 5’-TTCCTGGTGCACAACGTG‐ 3’, RP49-Rv, 5’-TCTCCTTGCGCTTCTTGG3’, Drosomycin-Fw, 5’-GTACTTGTTCGCCCTCTTCG‐ 3’, Drosomycin-Rv, 5’-CAGGGACCCTTGTATCTTCC‐ 3’, Defensin-Fw, 5’-CTTCGTTCTCGTGGCTATCG-3’, Defensin-Rv, 5’‐ CCAGGACATGATCCTCTGGA-3’, Diptericin-Fw, 5’-CAGTCCAGGGTCACCAGAAG-3’, Diptericin-Rv, 5’-AGGTGCTTCCCACTTTCCAG-3’, Metchnikowin-Fw, 5’-TACATCAGTGCTGGCAGAGC-3’, Metchnikowin-Rv, 5’-ACCCGGTCTTGGTTGGTTAG-3’, AttacinA-Fw, 5’-ACTACCTTGGATCTCACGGGA-3’, AttacinA-Rv, 5’‐ TGATGAGATAGACCCAGGCCA-3’, Cecropin-Fw, 5’-ATCGGAAGCTGGTTGGCTAAA-3’, Cecropin-Rv, 5’-GTGGTTAACCTCGAGCAGTGG-3’, TurandotB-Fw, 5’-CGCATGGCTCCTAGCTTAAGA‐ 3’, TurandotB-Rv, 5’-CTGGGTACTCCATCGACCATG‐ 3’. Each sample was duplicated on the PCR plate, and the final results average three biological replicates. For the quantification, the ΔΔCt method was used to determine the differences between target gene expression relative to the reference *Rp49* gene expression.

### Lymph gland immunostaining

For immunostaining of larval lymph glands, LGs were dissected from matured 3rd instar larvae and fixed in 4.0 % paraformaldehyde in PBS for 15 min at 25 °C. After repeated washing, samples were blocked with PBS containing 0.1 % Triton X-100 and 10 % normal goat serum and the fixed samples were incubated with primary antibodies at 4 °C for overnight. The following antibodies were used as primary antibodies: anti-DE-cadherin monoclonal antibody (diluted at 1: 300; DSHB: Cat&DCAD2-c) and anti-HA-tag rabbit IgG (1: 200; Cell Signaling Technology, MA, USA). After extensive washing, specimens were incubated with Alexa 594 or Alexa 488 secondary antibodies (1: 400; Molecular Probe, CA, USA). The LG specimens were placed on a fluorescence microscope (Olympus, Tokyo, Japan, model: IX81), outfitted with excitation, emission filter wheels (Olympus). The fluorescence signals were collected using a 10x, 20x,40x or 60x dry objective lens. Specimens were illuminated with UV filtered and shuttered light using the appropriate filter wheel combinations through a GFP/RFP filter cube. Near simultaneous GFP and/or RFP fluorescence images were captured with a CCD camera (Hamamatsu Photonics, Shizuoka, Japan). Image acquisition was controlled through the Metamorph software version 7.6 (Molecular Devices, Sunnyvale, CA, USA) and processed with ImageJ or Adobe Photoshop CS.

### Immunostaining of circulating hemocytes

After washing of 3rd instar larvae in PBS, single larva was transferred into *Drosophila* Ringer solution (DR) (10 mM, pH 7.2 Tris-HCl, 3 mM CaCl_2_ · 2H_2_O, 182 mM KCL, 46 mM NaCl) on a slide glass. And then, only the larval epidermis was cut by a set of fine forceps so as to allow circulating hemocytes to release into the DR outside the larvae. After an aliquot of DR containing circulating hemocytes was placed on the slide glass to evaporate it by a hot air, hemocytes were fixed in 4.0 % paraformaldehyde for 5 min at 25 °C. Immunostaining of hemocytes was carried out as described above.

### Apoptosis assay

Cells undergoing apoptosis were detected using a Cell Event Caspase-3/7 Green Detection Reagent (Molecular Probes, Invitrogen, CA, USA). 3rd instar larvae were dissected in PBS to collect LGs or imaginal discs. They were, then incubated in PBS containing 2 μM Cell Event Caspase3/7 Green Detection Reagent for 30 min at 37 °C. After incubation, the LGs or imaginal discs were fixed with 4.0 % paraformaldehyde for 15 min at 25 °C. After a fixation, tissue specimens were repeatedly washed and permeabilized in PBST. After several washings, specimens were mounted and observed as described above.

### Statistics

Results were presented as bar graphs or scatter plots created using GraphPad Prism 6. Every single set of data was assessed using Student’s *t*-test. Statistical significance is described in each figure legend as follows; \**p* < 0.05, \*\**p* < 0.01, \*\*\**p* < 0.001 and \*\*\*\**p* < 0.0001.

## Acknowledgements

We thank Dr. N. Perrimon (Harvard Univ.), Dr. D. Hultmark (Umea Univ.), Dr. L. Bruno (Global Health Inst.), Dr. Y. Yagi (Nagoya Univ), Bloomington Stock Center, Fly ORF center and Drosophila Genetic Resource Center for providing the fly stocks.

## Data availability

RNA-seq data have been submitted to the Gene Expression Omnibus database (accession number: GSE121551).

## Author contributions statement

MA and RA carried out observations of larval lymph glands, immunostaining experiments of the larval tissues and the qRT-PCR experiments. YO performed RNA-sequencing. TS analysed the RNA-seq data. YHI planned, organized the project, led the interpretation of the data, wrote, and revised the manuscript. All authors read and approved the final manuscript.

## References

An, S., Dong, S., Wang, Q., Li, S., Gilbert, L.I., Stanley, D., and Song, Q. (2012). Insect neuropeptide bursicon homodimers induce innate immune and stress genes during molting by activating the NF‐κB transcription factor Relish. PLOS ONE 7, e34510.

Arbouzova, N.I., and Zeidler, M.P. (2006). JAK/STAT signalling in Drosophila: insights into conserved regulatory and cellular functions. Development 133, 2605–2616.

Baker, M.A., Maloy, W.L., Zasloff, M., and Jacob, L.S. (1993). Anticancer Efficacy of Magainin2 and Analogue Peptides. Cancer Res. 53, 3052–3057.

Belvin, M.P., and Anderson, K.V. (1996). A conserved signaling pathway: The Drosophila Toll-Dorsal pathway. Annu. Rev. Cell Dev. Biol. 12, 393–416.

Brennan, C.A., and Anderson, K.V. (2004). Drosophila: The genetics of innate immune recognition and response. Annu. Rev. Immunol. 22, 457–483.

Choe, K.-M., Werner, T., Stöven, S., Hultmark, D., and Anderson, K.V. (2002). Requirement for a peptidoglycan recognition protein (PGRP) in relish activation and antibacterial immune responses in Drosophila. Science 296, 359–362.

Choe, K.-M., Lee, H., and Anderson, K.V. (2005). Drosophila peptidoglycan recognition protein LC (PGRP-LC) acts as a signal-transducing innate immune receptor. Proc. Natl. Acad. Sci. U. S. A. 102, 1122–1126.

Cordero, J.B., Macagno, J.P., Stefanatos, R.K., Strathdee, K.E., Cagan, R.L., and Vidal, M. (2010). Oncogenic Ras diverts a host TNF tumor suppressor activity into tumor promoter. Dev. Cell 18, 999–1011.

Destoumieux, D., Munoz, M., Bulet, P., and Bachère, E. (2000). Penaeidins, a family of antimicrobial peptides from penaeid shrimp (Crustacea, Decapoda). Cell. Mol. Life Sci. CMLS 57, 1260–1271.

Domazet-Lošo, T., Klimovich, A., Anokhin, B., Anton-Erxleben, F., Hamm, M.J., Lange, C., and Bosch, T.C.G. (2014). Naturally occurring tumours in the basal metazoan Hydra. Nat. Commun. 5, 4222.

Dostert, C., Jouanguy, E., Irving, P., Troxler, L., Galiana-Arnoux, D., Hetru, C., Hoffmann, J.A., and Imler, J.-L. (2005). The Jak-STAT signaling pathway is required but not sufficient for the antiviral response of Drosophila. Nat. Immunol. 6, 946.

Ertürk-Hasdemir, D., Broemer, M., Leulier, F., Lane, W.S., Paquette, N., Hwang, D., Kim, C.-H., Stöven, S., Meier, P., and Silverman, N. (2009). Two roles for the Drosophila IKK complex in the activation of Relish and the induction of antimicrobial peptide genes. Proc. Natl. Acad. Sci. 106, 9779–9784.

Fehlbaum, P., Bulet, P., Michaut, L., Lagueux, M., Broekaert, W.F., Hetru, C., and Hoffmann, J.A. (1994). Insect immunity. Septic injury of Drosophila induces the synthesis of a potent antifungal peptide with sequence homology to plant antifungal peptides. J. Biol. Chem. 269, 33159–33163.

Gaspar, D., Veiga, A.S., and Castanho, M.A.R.B. (2013). From antimicrobial to anticancer peptides. A review. Front. Microbiol. 4.

Gottar, M., Gobert, V., Michel, T., Belvin, M., Duyk, G., Hoffmann, J.A., Ferrandon, D., and Royet, J. (2002). The Drosophila immune response against gram-negative bacteria is mediated by a peptidoglycan recognition protein. Nature 416, 640.

Gray, B. (2011). PSVue®: Novel small molecule probes for fluorescent imaging of apoptosis and bacterial infections in vitro and in vivo. Nat. Methods Appl. Notes.

Hoffmann, J.A. (2003). The immune response of Drosophila. Nature 426, 33.

Hoffmann, J.A., and Reichhart, J.-M. (2002). Drosophila innate immunity: an evolutionary perspective. Nat. Immunol. 3, 121.

Hoskin, D.W., and Ramamoorthy, A. (2008). Studies on anticancer activities of antimicrobial peptides. Biochim. Biophys. Acta BBA - Biomembr. 1778, 357–375.

Jang, I.-H., Chosa, N., Kim, S.-H., Nam, H.-J., Lemaitre, B., Ochiai, M., Kambris, Z., Brun, S., Hashimoto, C., Ashida, M., et al. (2006). A Spätzle-processing enzyme required for Toll signaling activation in Drosophila innate immunity. Dev. Cell 10, 45–55.

Kim, R., Emi, M., and Tanabe, K. (2007). Cancer immunoediting from immune surveillance to immune escape. Immunology. 121, 1-14.

Kim, D., Pertea, G., Trapnell, C., Pimentel, H., Kelley, R., and Salzberg, S.L. (2013). TopHat2: accurate alignment of transcriptomes in the presence of insertions, deletions and gene fusions. Genome Biol. 14, R36.

Landais, S., D’Alterio, C., and Jones, D.L. (2014). Persistent replicative stress alters polycomb phenotypes and tissue homeostasis in Drosophila melanogaster. Cell Rep. 7, 859–870.

Lanot, R., Zachary, D., Holder, F., and Meister, M. (2001). Postembryonic hematopoiesis in Drosophila. Dev. Biol. 230, 243–257.

Lemaitre, B., and Hoffmann, J. (2007). The host defense of Drosophila melanogaster. Annu. Rev. Immunol. 25, 697–743.

Lemaitre, B., Reichhart, J.-M., and Hoffmann, J.A. (1997). Drosophila host defense: Differential induction of antimicrobial peptide genes after infection by various classes of microorganisms. Proc. Natl. Acad. Sci. U. S. A. 94, 14614–14619.

Lim, J.-H., Kim, M.-S., Kim, H.-E., Yano, T., Oshima, Y., Aggarwal, K., Goldman, W.E., Silverman, N., Kurata, S., and Oh, B.-H. (2006). Structural basis for preferential recognition of diaminopimelic acid-type peptidoglycan by a subset of peptidoglycan recognition proteins. J. Biol. Chem. 281, 8286–8295.

Mader, J.S., and Hoskin, D.W. (2006). Cationic antimicrobial peptides as novel cytotoxic agents for cancer treatment. Expert Opin. Investig. Drugs 15, 933–946.

Morisato, D., and Anderson, K.V. (1994). The spätzle gene encodes a component of the extracellular signaling pathway establishing the dorsal-ventral pattern of the Drosophila embryo. Cell 76, 677–688.

Oka, S., Hirai, J., Yasukawa, T., Nakahara, Y., and Inoue, Y.H. (2015). A correlation of reactive oxygen species accumulation by depletion of superoxide dismutases with age-dependent impairment in the nervous system and muscles of Drosophila adults. Biogerontol. 16, 485–501.

Papo, N., and Shai, Y. (2005). Host defense peptides as new weapons in cancer treatment. Cell. Mol. Life Sci. CMLS 62, 784–790.

Parisi, F., Stefanatos, R.K., Strathdee, K., Yu, Y., and Vidal, M. (2014). Transformed epithelia trigger non-tissue-autonomous tumor suppressor response by adipocytes via activation of Toll and Eiger/TNF signaling. Cell Rep. 6, 855–867.

Pastor-Pareja, J.C., Wu, M., and Xu, T. (2008). An innate immune response of blood cells to tumors and tissue damage in Drosophila. Dis. Model. Mech. 1, 144–154.

Porter, A.G., and Jänicke, R.U. (1999). Emerging roles of caspase-3 in apoptosis. Cell Death Differ. 6, 99.

Rawlings, J.S., Rosler, K.M., and Harrison, D.A. (2004). The JAK/STAT signaling pathway. J. Cell Sci. 117, 1281–1283.

Remillieux-Leschelle, N., Santamaria, P., and Randsholt, N.B. (2002). Regulation of Larval Hematopoiesis in Drosophila melanogaster: A Role for the multi sex combs Gene. Genetics 162, 1259–1274.

Rivero-Muller, A., Vuorenoja, S., Tuominen, Wacławik, A., Brokken, LJS., Ziecik, A.J., Huhtaniemi, I., Rahman, N.A. (2007) Use of hecate–chorionic gonadotropin β conjugate in therapy of lutenizing hormone receptor expressing gonadal somatic cell tumors. Mol. Cell. Endocrinol. 269, 17-25.

Rizki, T.M., and Rizki, R.M. (1980). Properties of the larval hemocytes of Drosophila melanogaster. Experientia 36, 1223–1226.

Saget, O., Forquignon, F., Santamaria, P., and Randsholt, N.B. (1998). Needs and targets for the multi sex combs gene products in Drosophila melanogaster. Genetics 149, 1823-1838.

Santamaria, P., and Randsholt, N.B. (1995). Characterization of a region of the X chromosome of Drosophila including multi sex combs (mxc), a Polycomb group gene which also functions as a tumour suppressor. Mol. Gen. Genet. MGG 246, 282–290.

Shrestha, R., Gateff, E. (1982). Ultrastructure and cytochemistry of the cell types in the larval hematopoietic organs and haemolymph of Drosophila melanogaster. Dev. Growth Differ. 24, 65-82.

Sloballe, P.W., Lee Maloy, W., Myrga, M.L., Jacob, L.S., and Herlyn, M. (1995). Experimental local therapy of human melanoma with lytic magainin peptides. Int. J. Cancer 60, 280–284.

Stöven, S., Ando, I., Kadalayil, L., Engström, Y., and Hultmark, D. (2000). Activation of the Drosophila NF‐ _κ_ B factor Relish by rapid endoproteolytic cleavage. EMBO Rep. 1, 347–352.

Takehana, A., Yano, T., Mita, S., Kotani, A., Oshima, Y., and Kurata, S. (2004). Peptidoglycan recognition protein (PGRP)‐LE and PGRP‐LC act synergistically in Drosophila immunity. EMBO J. 23, 4690–4700.

Tanji, T., and Ip, Y.T. (2005). Regulators of the Toll and Imd pathways in the Drosophila innate immune response. Trends Immunol. 26, 193–198.

Trapnell, C., Roberts, A., Goff, L., Pertea, G., Kim, D., Kelley, D.R., Pimentel, H., Salzberg, S.L., Rinn, J.L., and Pachter, L. (2012). Differential gene and transcript expression analysis of RNA-seq experiments with TopHat and Cufflinks. Nat. Protoc. 7, 562–578.

Tzou, P., De Gregorio, E., and Lemaitre, B. (2002). How Drosophila combats microbial infection: a model to study innate immunity and host-pathogen interactions. Curr Opin Microbiol 5, 102–110.

Valanne, S., Wang, J.-H., and Rämet, M. (2011). The Drosophila Toll Signaling Pathway. J. Immunol. 186, 649–656.

Winder, D., Günzburg, W.H., Erfle, V., and Salmons, B. (1998). Expression of antimicrobial peptides has an antitumour effect in human cells. Biochem. Biophys. Res. Commun. 242, 608–612.

Yakovlev, A.Y., Nesin, A.P., Simonenko, N.P., Gordya, N.A., Tulin, D.V., Kruglikova, A.A., and Chernysh, S.I. (2017). Fat body and hemocyte contribution to the antimicrobial peptide synthesis in Calliphora vicina R.-D. (Diptera: Calliphoridae) larvae. Vitro Cell. Dev. Biol. - Anim. 53, 33–42.

Zaiou, M. (2007). Multifunctional antimicrobial peptides: therapeutic targets in several human diseases. J. Mol. Med. 85, 317–329.

Zhang, L., and Gallo, R.L. (2016). Antimicrobial peptides. Curr. Biol. 26, R14–R19.

Zou, H., Li, Y., Liu, X., and Wang, X. (1999). An APAF-1•cytochrome c multimeric complex is a functional apoptosome that activates procaspase-9. J. Biol. Chem. 274, 11549–11556.

